# Identification of novel tail-anchored membrane proteins integrated by the bacterial twin-arginine translocase

**DOI:** 10.1101/2023.11.02.564894

**Authors:** José Jesús Gallego-Parrilla, Emmanuele Severi, Govind Chandra, Tracy Palmer

## Abstract

The twin arginine protein transport (Tat) system exports folded proteins across the cytoplasmic membranes of prokaryotes and the energy transducing membranes of plant thylakoids and mitochondria. Proteins are targeted to the Tat machinery by N-terminal signal peptides with a conserved twin arginine motif, and some substrates are exported as heterodimers where the signal peptide is present on one of the partner proteins. A subset of Tat substrates is found in the membrane. Tat-dependent membrane proteins usually have large globular domains and a single transmembrane helix present at the N- or C-terminus. Five Tat substrates that have C-terminal transmembrane helices have previously been characterised in the model bacterium *Escherichia coli*. Each of these is an iron-sulphur cluster-containing protein involved in electron transfer from hydrogen or formate. Here we have undertaken a bioinformatic search to identify further tail-anchored Tat substrates encoded in bacterial genomes. Our analysis has revealed additional tail-anchored iron-sulphur proteins associated in modules with either a *b*-type cytochrome or a quinol oxidase. We also identified further candidate tail-anchored Tat substrates, particularly among members of the actinobacterial genus, that are not predicted to contain cofactors. Using reporter assays we show experimentally that six of these have both N-terminal Tat signal peptides and C-terminal transmembrane helices. The newly-identified proteins include a carboxypeptidase and a predicted protease, and four sortase substrates for which membrane integration is a pre-requisite for covalent attachment to the cell wall.

## INTRODUCTION

Membrane proteins play critical roles in all cells, and in bacteria cytoplasmic membrane proteins comprise approximately 20 – 30% of the proteome (1, 2). Integral membrane proteins vary in length, hydrophobicity and overall topology, with some containing a single transmembrane segment (TMH) whereas others span the membrane multiple times. Polytopic membrane proteins are usually synthesised and inserted into the membrane co-translationally, to prevent cytoplasmic aggregation. This is achieved through recognition of the first hydrophobic stretch of the nascent protein by ribosome-associated signal recognition particle (SRP). The translating ribosome is guided to the membrane through interaction of SRP with its receptor, and the polypeptide is threaded through the Sec translocon with hydrophobic helices exiting through the lateral gate into the lipid bilayer. By contrast, some smaller membrane proteins, usually comprising one or two TMH with short loop regions, are integrated Sec-independently by the membrane insertase YidC (reviewed in (3, 4)).

Alongside the Sec pathway and YidC, the twin arginine translocase (Tat) is the third system found in the cytoplasmic membrane of prokaryotes that is able to mediate membrane protein insertion. Although most Tat substrates identified to date are globular periplasmic proteins, the Tat pathway is able to integrate proteins that have a single hydrophobic helix located either at the N- or C-terminus of the substrate (5, 6), or occasionally within the protein sequence (7, 8).

The major distinguishing feature of the Tat system is that it transports fully folded proteins (9). Tat substrates are targeted for export by N-terminal signal peptides that harbour a twin arginine motif. The motif is minimally defined as R-R-x-ϕ-ϕ where x is any amino acid and ϕ represents a hydrophobic amino acid (10–12). The twin arginines are almost always invariant and are critical for recognition by the Tat machinery (13, 14). Tat signal peptides have a tripartite arrangement with the twin arginines conferring a positive charge to the n-region.

This precedes a hydrophobic h-region and a polar c-region that contains a cleavage site for signal peptidase (10, 15). Sec signal peptides are very similar to those that target the Tat pathway but generally have less sequence constraints. For example, the n-region positive charge of a Sec signal may be conferred by a single arginine or lysine, or by two or more of these side chains (16). Sec and Tat signal peptides also differ in the hydrophobicity of their respective h-regions; Tat signal peptides are significantly less hydrophobic, and this property avoids mistargeting of Tat substrates to the Sec pathway rather than being mechanistically essential for Tat transport (17, 18). Likewise, the presence of one or more positive charges in Tat signal peptide c-regions and/or in the first few amino acids of the mature domain also serves to avoid productive engagement with the Sec pathway (19, 20). Nonetheless, despite these differences, there is overlap between the two classes of targeting sequences, and some twin-arginine signal peptides from *Escherichia coli* Tat substrates can mediate export by the Sec pathway if they are fused to a compatible reporter protein (20).

Many Tat substrates contain redox cofactors such as iron-sulphur clusters or molybdopterins and are components of electron transport chains (21, 22). Cofactor insertion into the apoprotein is catalysed in the cytoplasm, which results in irreversible folding, necessitating export by the Tat pathway (10). Some Tat substrates, for example the periplasmic Ni-Fe hydrogenases and formate dehydrogenases are exported as heterodimers with the signal peptide being present on only one of the subunits (23, 24). The untagged subunits are therefore exported across the membrane solely because they form a complex with a partner bearing a twin-arginine signal peptide in a ‘hitchhiker’ mechanism (23). A further interesting feature of these particular Tat substrates is that they are anchored to the periplasmic face of the inner membrane by a C-terminal TMH (‘C-tail’) present on one of the subunits (Fig. 1A). In the case of hydrogenase-1 (HYD-1) and hydrogenase-2 (HYD-2), the small subunit (HyaA for HYD-1 and HybO for HYD-2) bears both the Tat signal peptide and the C-tail (5, 25). By contrast, the twin arginine signal peptide is found on the catalytic subunit of formate dehydrogenases whereas the C-tail is present on the smaller iron-sulphur partner (26). A further, tail-anchored Tat substrate, HybA, is a monomeric iron-sulphur protein that forms part of the HYD-2 electron transport pathway (5, 27) (Fig 1A).

**Fig 1.**
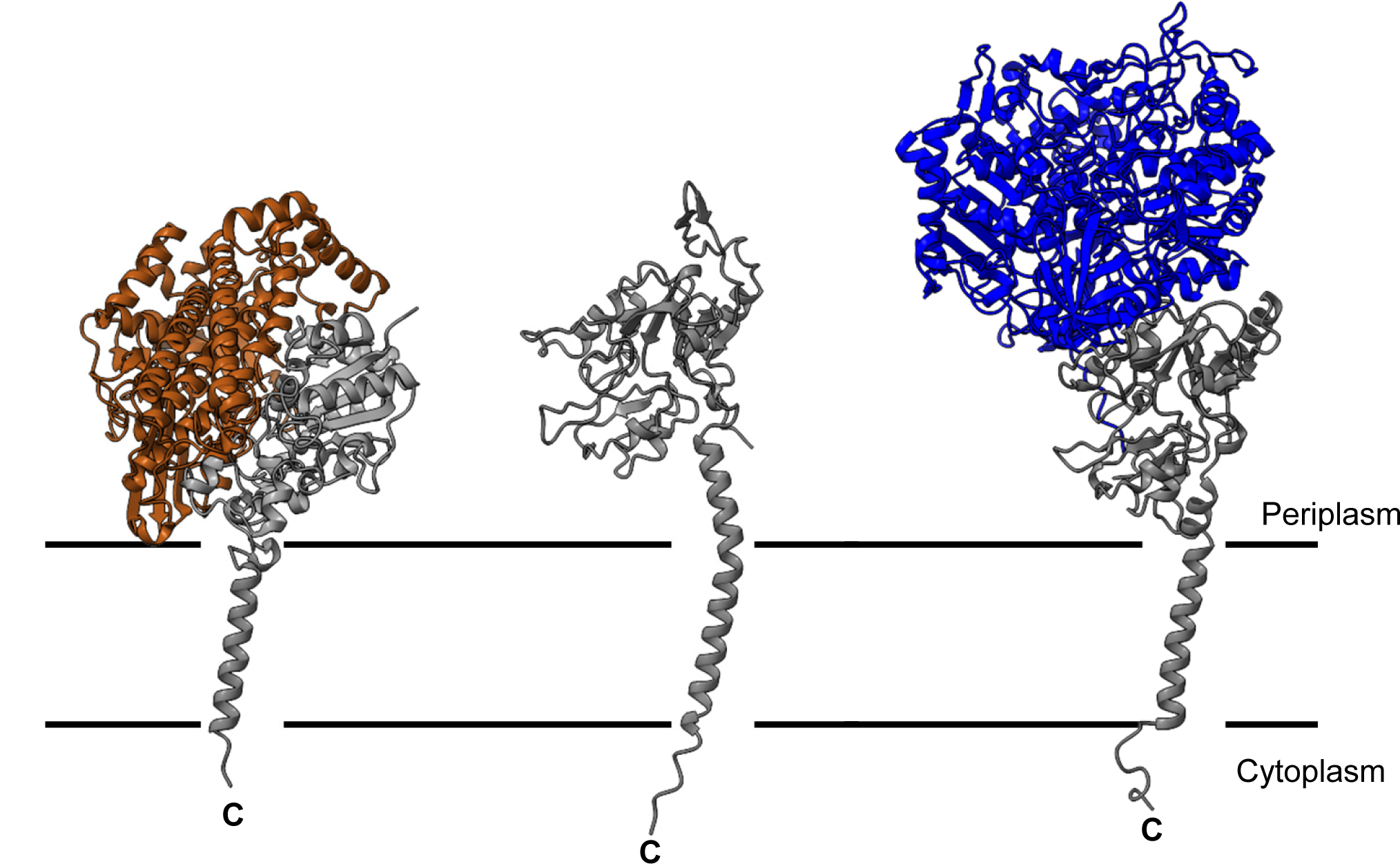
Structures of C-tail containing Tat substrates. Left – periplasmic Ni-Fe hydrogenase (*E. coli* HYD-1; pdb 4GD3); right periplasmic formate dehydrogenase (*E. coli* FDH-N; pdb 1KQF). Note that in each case the membrane-embedded cytochrome subunits have not been included. Centre – structural model of *E. coli* HybA generated using ESMFold (65). In each case the C-tail containing subunit is shown in grey. All metallocofactors have been omitted for clarity.

While three families of Tat-dependent C-tail proteins have been identified, it is likely that further families remain to be discovered. Here we describe a bioinformatic approach to identify Tat-dependent tail-anchored membrane proteins. We identify additional tail-anchored iron-sulphur proteins encoded in genetic modules with cytochromes or quinol oxidases. We also identify numerous candidate proteins that are not predicted to bind any cofactors. Experimental validation using reporter fusions confirms that six of these represent novel Tat-dependent C-tail proteins.

## METHODS

### Bioinformatic methods

89,292 bacterial genomes labelled as “reference” or “representative” in Genbank (September 2017) were searched for Tat substrates using TATFind version 1.4 (11, 28). TMHMM-2.0c (2) program licensed (free academic license) from DTU was subsequently used to predict TMH in each protein predicted to be a Tat substrate by TATFind 1.4. If the protein was longer than 150 amino acids, the number of TMH in the protein was not more than two and, one of the helices ended within the last 50 amino acids of the protein then it was considered a positive find. Operationally, the TATFind 1.4 search, the TMHMM-2.0c search and, the filtering were all wrapped in a bespoke Perl script relying on BioPerl libraries (https://bioperl.org/). GNU Parallel was used to parallelise the searches (to speed up searching of over 89k genomes). PostgreSQL (https://www.postgresql.org/) was used for storage and management of search results.

The wanda.py script was designed to retrieve and process protein information from the NCBI database. The script takes a list of protein accession numbers as input and uses the Entrez module from the BioPython library to fetch detailed information for each protein from NCBI’s Entrez service. The retrieved information includes the protein ID, source, nucleotide accession, start and stop positions, strand, protein name, organism, strain, and assembly. This information is written into a tab-separated value (TSV) file named ‘efetch_output.tsv’. The script then processes this file using the pandas library, removing rows sourced from ‘INSDC’ and rearranging the columns for easier viewing. It also sorts the data based on the assembly number and removes any duplicates based on a subset of fields (start, stop, strand, organism, strain, assembly). Finally, the script sorts the processed data alphabetically by organism and writes it to a new TSV file named ‘final_parsed_output.tsv’. The script wanda.py is available in our GitHub repository (https://github.com/Ravenneo/Ctails_Tat_Proteins).

Genomic neighbourhoods flanking genes of interest were analysed using FlaGs (29) and visualised with Clinker (30). Proteins homologous to proteins of interested were identified using BLAST-P (https://blast.ncbi.nlm.nih.gov/Blast.cgi). The presence of signal peptides in proteins was predicted using SignalP-6.0 (31) or through DeepTMHMM for the presence of transmembrane regions (32).

### Strains and plasmids

All *E. coli* strains used throughout this study are isogenic derivatives of MC4100 (33). Protein fusions between SufI and candidate C-tails (“CTs”) were expressed and characterised in strain NRS-3 (As, MC4100 Δ*sufI*; (34)), while protein fusions between the mature sequence of AmiA and test signal peptides (“SPs”) were studied in strains MC4100 Δ*amiA* Δ*amiC* and MC4100 Δ*amiA* Δ*amiC* Δ*tatABC* (35). Strains XL1blue (Agilent), XL10-Gold (Agilent), and DH5α (ThermoFisher) were used for cloning.

All plasmid constructs and oligonucleotides used in this study are listed in Tables 1 and S1, respectively. Q5 (NEB) and Verify (PCRBIO) were used for high-fidelity PCR. To construct *amiA* and *sufI* gene fusions, the coding sequences (CDS) of target SP and CT peptides were commercially synthesised (IDT, GenScript) as fragments codon-optimised for expression in *E. coli* (sequences available in Table S2). These were then used as templates for PCR reactions to add suitable restriction sites for restriction-ligation cloning into the appropriate recipient plasmids. For SP-*amiA* cloning, SP-coding PCR products were digested with *Xba*I and *Bam*HI and ligated between the same sites of pSU40UniAmiA (36). For *sufI*-CT cloning, we first constructed plasmid pESN337 bearing the *sufI*-CT*_fdnH_*fusion (with the fusion reproducing the design of Hatzixanthis *et al*. (5)) by three-way ligation of pSUPROM (37) digested with *Bam*HI and *Sph*I, a *sufI* PCR product (oligos JES3+JES4 amplified from MG1655 gDNA) digested with *Bam*HI and *Sac*I, and a PCR product covering the 3’-end of *fdnH* (oligos ESN584+ESN585 amplified from MG1655 gDNA) digested with *Sac*I and *Sph*I.

**Table 1:** Plasmids used and constructed in this study. The method of construction of each plasmid is indicated in in parentheses together with the names of primer pairs used to amplify PCR products (primer sequences listed in Table S1). Methods are abbreviated as follows: RL, restriction-ligation; KLD: whole-plasmid mutagenic PCR circularised with NEB’s KLD kit; QC: QuickChange mutagenesis. SP: signal peptide (CDS); CT: C-tail (CDS). * indicates a longer version of C-tail 10 from a different protein accession.

| PLASMID | DESCRIPTION | REFERENCE |
| --- | --- | --- |
| pSU-PROM | *tatA* promoter + MCS in pSU40 backbone. Kan<sup>R</sup> | (37) |
| pAmiA | pSU-PROM encoding AmiA lacking its native signal peptide | (36) |
| pTorASP-AmiA | As pAmiA with TorA signal peptide cloned in frame with AmiA. | (36) |
| pJGP1 | As pAmiA with SP1 cloned in frame with AmiA (RL SP1F+SP1R) | This work |
| pJGP2 | As pAmiA with SP2 cloned in frame with AmiA (RL SP2F+SP2R) | This work |
| pESN328 | As pAmiA with SP3 cloned in frame with AmiA (RL ESN540+ESN541) | This work |
| pJGP3 | As pAmiA with SP4 cloned in frame with AmiA (RL SP4F+SP4R) | This work |
| pESN329 | As pAmiA with SP5 cloned in frame with AmiA (RL ESN542+ESN543) | This work |
| pESN330 | As pAmiA with SP6 cloned in frame with AmiA (RL ESN544+ESN545) | This work |
| pESN331 | As pAmiA with SP7 cloned in frame with AmiA (RL ESN546+ESN547) | This work |
| pJGP4 | As pAmiA with SP8 cloned in frame with AmiA (RL SP8F+SP8R) | This work |
| pJGP5 | As pAmiA with SP9 cloned in frame with AmiA (RL SP9F+SP9R) | This work |
| pESN355 | As pAmiA with SP9-[C24A] cloned in frame with AmiA (KLD ESN675+ESN676) | This work |
| pESN361 | As pAmiA with SP10 cloned in frame with AmiA (RL ESN102+ESN434) | This work |
| pESN362 | As pAmiA with SP11 cloned in frame with AmiA (RL ESN102+ESN434) | This work |
| pESN363 | As pAmiA with SP12 cloned in frame with AmiA (RL ESN102+ESN434) | This work |
| pESN364 | As pAmiA with SP13 cloned in frame with AmiA (RL ESN102+ESN434) | This work |
| pESN365 | As pAmiA with SP14 cloned in frame with AmiA (RL ESN102+ESN434) | This work |
| pESN366 | As pAmiA with SP15 cloned in frame with AmiA (RL ESN102+ESN434) | This work |
| pESN367 | As pAmiA with SP16 cloned in frame with AmiA (RL ESN102+ESN434) | This work |
| pESN339 | As pSU-PROM encoding native *sufI* (QC ESN591+ESN592) | This work |
| pESN337 | pSU-PROM encoding *SufI*-CT<sub>fdnH</sub> (RL JES3+JES4 and ESN584+ESN585) | This work |
| pJGP6 | pSU-PROM encoding *SufI*-CT1 (RL CT1F+CT1R) | This work |
| pJGP7 | pSU-PROM encoding *SufI*-CT2 (RL CT2F+CT2R) | This work |
| pESN332 | pSU-PROM encoding *SufI*-CT3 (RL ESN550+ESN551) | This work |
| pESN333 | pSU-PROM encoding *SufI*-CT4 (RL ESN552+ESN553) | This work |
| pESN334 | pSU-PROM encoding *SufI*-CT5 (RL ESN554+ESN555) | This work |
| pJGP8 | pSU-PROM encoding *SufI*-CT6 (RL CT6F+CT6R) | This work |
| pESN335 | pSU-PROM encoding *SufI*-CT7 (RL ESN556+ESN557) | This work |
| pJGP9 | pSU-PROM encoding *SufI*-CT8 (RL CT8F+CT8R) | This work |
| pESN336 | pSU-PROM encoding *SufI*-CT9 (ESN565+ESN566) | This work |
| pESN368 | pSU-PROM encoding Sufl-CT10 (ESN103+ESN237) | This work |
| pESN376 | pSU-PROM encoding Sufl-CT10* (ESN103+ESN237) | This work |
| pESN369 | pSU-PROM encoding Sufl-CT11 (ESN103+ESN237) | This work |
| pESN370 | pSU-PROM encoding Sufl-CT12 (ESN103+ESN237) | This work |
| pESN371 | pSU-PROM encoding Sufl-CT13 (ESN103+ESN237) | This work |
| pESN372 | pSU-PROM encoding Sufl-CT14 (ESN103+ESN237) | This work |
| pESN373 | pSU-PROM encoding Sufl-CT15 (ESN103+ESN237) | This work |
| pESN374 | pSU-PROM encoding Sufl-CT16 (ESN103+ESN237) | This work |

All other *sufI*-CT fusion constructs were derived from pESN337 by subcloning CT-coding PCR products between the *Sac*I and *Sph*I sites of the construct, replacing the coding region of CT*_fdnH_*. All mutant derivatives of these plasmids were made by whole-plasmid PCR-based techniques, either by QuickChange (38) or using NEB’s KLD kit, using appropriate parental plasmids as templates. All constructs were verified by sequencing.

### Growth experiments

Single colonies of freshly made transformants of MC4100 Δ*amiA* Δ*amiC* (± Δ*tatABC*) carrying pSU40UniAmiA-based constructs were grown overnight in LB with 50 mgml^-1^ kanamycin (LBKan). These were diluted to an initial OD_600_ of 0.05 in LBKan with or without 0.5% SDS (sodium dodecyl sulphate) and grown in a TECAN model plate reader at 37 °C without shaking for 20 h with sampling every 20 min. Growth data were processed in Excel.

### Cell fractionation and Western Blotting

Single colonies of freshly made NRS-3 transformants carrying pSU-PROM-*sufI*-CT constructs were grown overnight in LBKan. These were diluted 1:100 in the same medium and grown until OD_600_ reached 0.3, at which point cells were harvested from 25 ml culture for cellular fractionation. Fractionation was performed as described in Severi *et al*. (39) with the addition of a urea wash step adapted from Hatzixanthis *et al*. (5). Briefly, cells were resuspended in 1 ml of ResB+EDTA buffer (20 mM Tris-Cl, 200 mM NaCl, 12 mM EDTA, pH 7.5) and ruptured by sonication. After clarification, the membrane fraction was separated by ultracentrifugation. While the top 200 ml of the supernatant were retained as the soluble fraction (cytoplasm + periplasm), the membrane pellets were resuspended in 1 ml ResB-EDTA supplemented with 8 M urea to remove loosely associated material, followed by another ultracentrifugation step and final resuspension in 60 ml Buffer 2 (50 mM Tris-HCl, 5 mM MgCl_2_, 10 % v/v glycerol, pH 7.5). Equal volumes of membrane and soluble fractions were run on SDS-PAGE gradient gels (BioRad) and analysed by Western Blotting using anti-SufI (40) and anti-rabbit (BioRad) antibodies. Blots were developed with ECL (BioRad).

## RESULTS

### Database searches to identify Tat-dependent C-tail proteins

We first took a bioinformatic approach to identify candidate Tat-dependent C-tail proteins. We searched bacterial genome sequences in Genbank for proteins predicted to have a Tat targeting sequence, no more than two TMH in total (to allow for signal anchor sequences) with one TMH within 50 amino acids of the C-terminus. It should be noted this approach will identify proteins where the Tat signal and the C-tail are on the same polypeptide (for example HybA and hydrogenase small subunit), but not will not directly identify hitchhiking proteins such as the small subunit of formate dehydrogenase. None-the-less, the search, which covered 89,292 bacterial genomes, identified 34,634 proteins from 20,558 individual genomes belonging to 6,798 distinct organisms. The output of this search can be viewed in dataset S1.

### Iron-sulphur proteins are found among tail-anchored protein candidates

By far the greatest number of proteins identified from this analysis were iron-sulphur proteins, in particular the small subunits of hydrogenases, and HybA-like proteins involved in hydrogenase electron transport pathways. In total, 26,858 of the individual entries featured ‘hydrogenase’ in their annotation. We did, however, also find some iron-sulphur proteins that appeared to be unlinked to hydrogenases. For example, searching the annotation for ‘ferredoxin’, identified 36 proteins, of which one was a duplicate and was removed from the analysis. The remaining 35 proteins are listed in Table S3. These tail-anchored ferredoxin proteins are generally encoded in one of two distinct genomic loci (Table S3, Fig.2, Fig.3). The first of these is a pairing with *b*-type cytochrome (Fig 2a). Further Blast analysis indicates that this two gene locus is found widely in bacterial genomes in both Gram-negative and Gram-positive bacteria (some examples are given in Fig 2b). There is a lack of genomic conservation in the neighbourhood of this two gene cluster, so it is not clear whether they function as a stand-alone module or part of an unidentified electron transport pathway. Interestingly, in *Desulfofundulus salinum* it was noted that they form an apparent operon with *tatAC* genes (Fig 2b).

**Fig 2.**
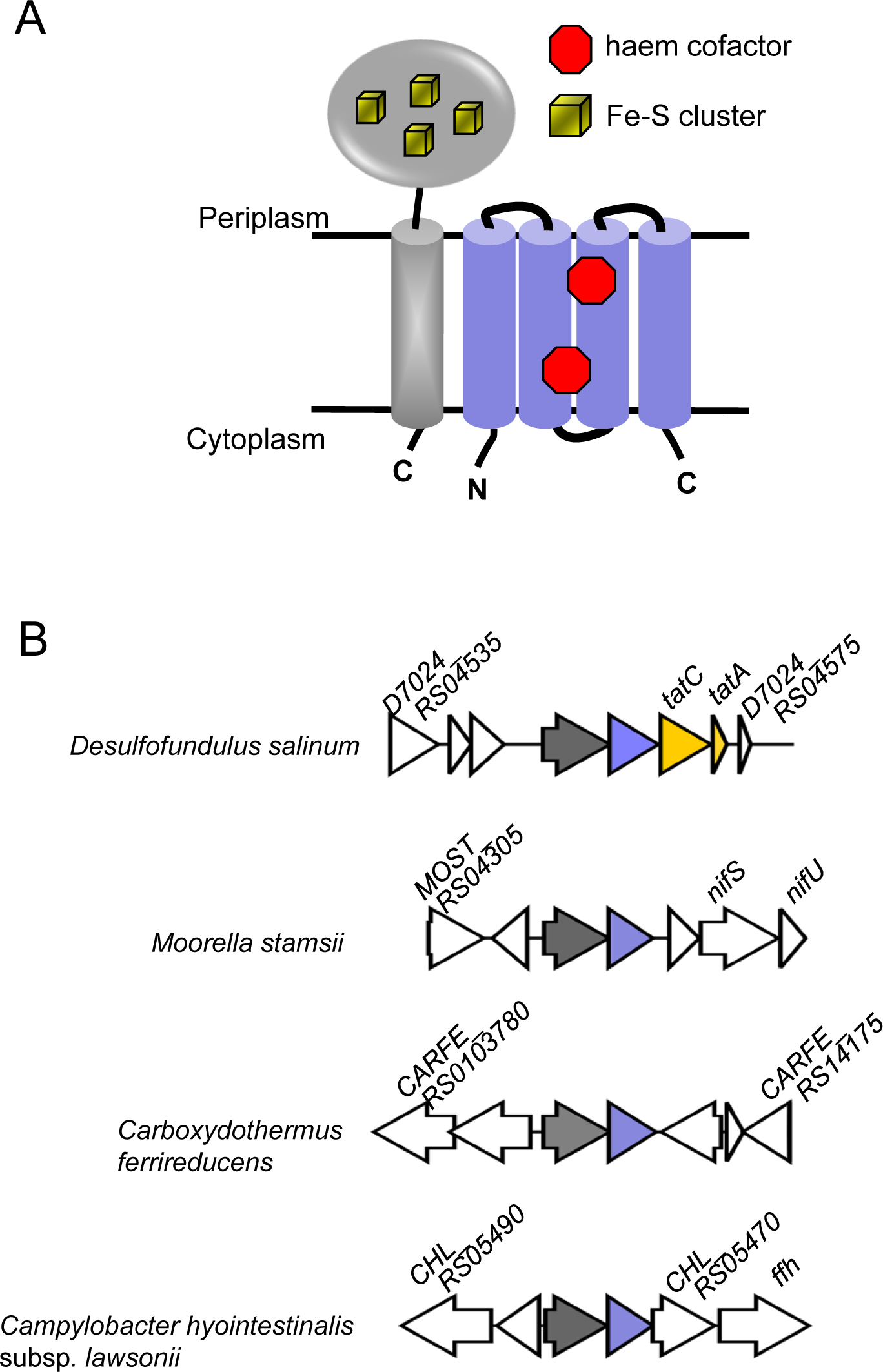
A C-tail containing iron-sulphur protein/cytochrome *b* module is encoded in many bacterial genomes. A. Predicted topologies of the Tat-dependent membrane anchored iron-sulphur protein (grey) and the FdnI-related cytochrome *b* (blue). B. Example loci encoding the two-protein module, with the iron-sulphur protein gene in grey and the cytochrome *b* gene in blue. Genes that are unrelated are shown in white. In *Desulfofundulus salinum* strain 435, *D7024_RS04535* is predicted to encode an ABC-type transporter, *D7024_RS04540* a YtrH family sporulation protein, *D7024_RS04545* a hypothetical protein and D7024_RS04575 a histidine kinase. In *Moorella stamsii* strain DSM 26217, *MOST_RS04305* is predicted to encode an Mrp/NBP35 family ATP-binding protein, *MOST_RS04310* a nitroreductase family protein and *MOST_RS04325* an Rrf2 family transcriptional regulator. In *Carboxydothermus ferrireducens* DSM 11255, *CARFE_RS0103780* is predicted to encode a class I SAM-dependent RNA methyltransferase, *CARFE_RS0103775* an MFS transporter, *CARFE_RS0103760* a radical SAM protein, *CARFE_RS16535* a 4Fe-4S binding protein and *CARFE_RS14175* a response regulator transcription factor pseudogene. In *Campylobacter hyointestinalis* subsp. *lawsonii*, *CHL_RS05490* is predicted to encode an FAD/NAD(P)-binding oxidoreductase, *CHL_RS05485* a Crp/Fnr family transcriptional regulator and *CHL_RS05470* a SLAC1 anion channel family protein.

**Fig 3.**
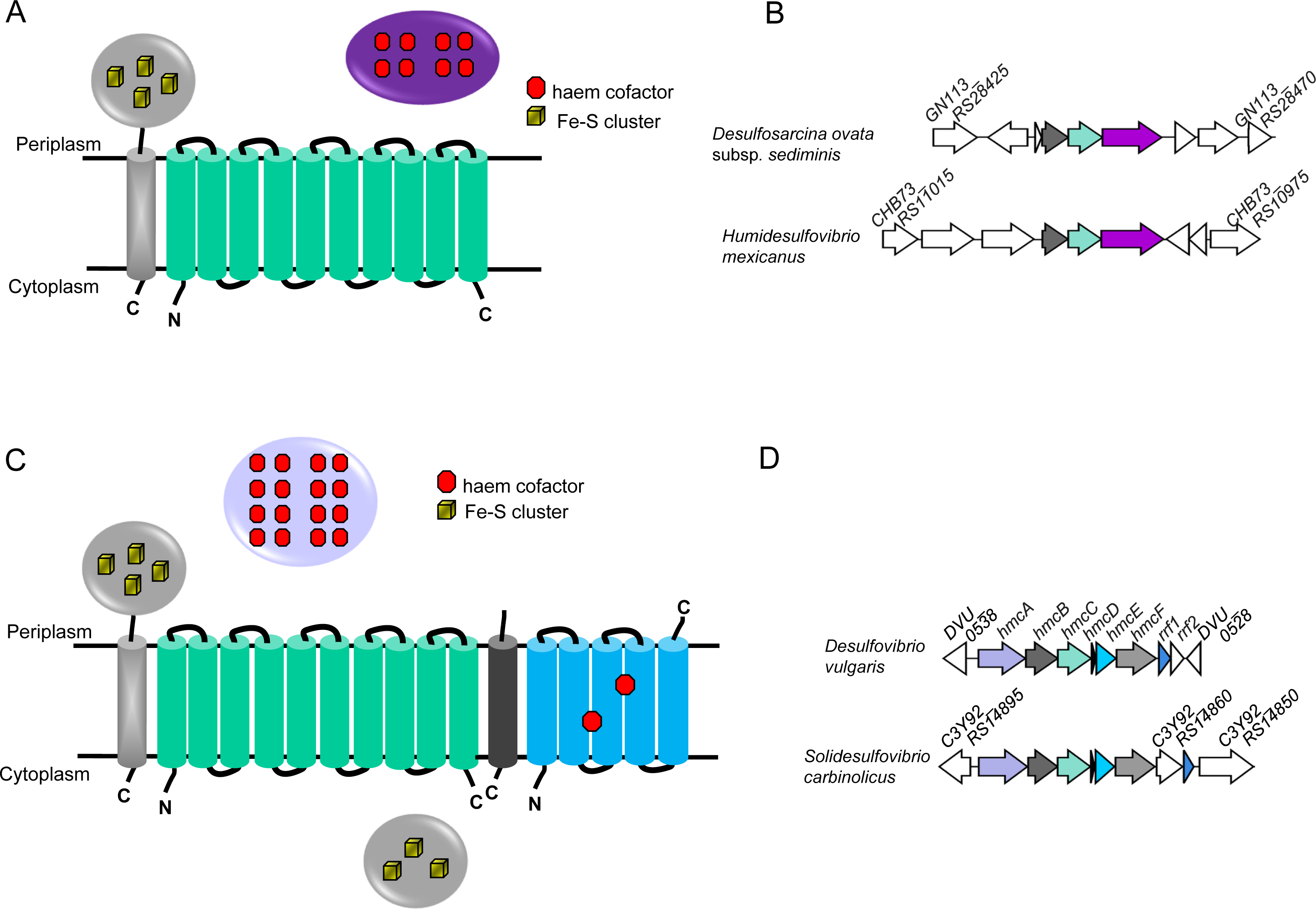
Co-occurrence of a C-tail iron-sulphur protein with a HybB-like quinol oxidase. A. Predicted topologies of the three-protein module comprising a C-tail containing iron-sulphur protein (grey), a HybB-like quinol oxidase lacking cofactors (cyan) and an octahaem *c*-type cytochrome (violet) encoded in bacterial genomes. B. Example loci encoding the three gene module, with the iron-sulphur protein gene in grey and the quinol oxidase gene in cyan and the cytochrome c gene in violet. Unrelated genes are shown in white. In *Desulfosarcina ovata* subsp. *sediminis* strain 28bB2T, *GN113_RS28425* is predicted to encode a YifB family Mg chelatase-like AAA ATPase, *GN113_RS28430* a sigma 54-interacting transcriptional regulator, *GN113_RS28435* a hypothetical protein, *prxU* a thioredoxin-dependent peroxiredoxin, *GN113_RS28465* an MBL fold metallo-hydrolase and *GN113_RS28470* an Mrp/NBP35 family ATP-binding protein. In *Humidesulfovibrio mexicanus* strain DSM 13116, *CHB73_RS11015* is predicted to encode an EAL-domain-containing protein, *CHB73_RS11010* a methyl-accepting chemotaxis protein, *CHB73_RS11005* a methyl-accepting chemotaxis protein, *cysQ* a 3’(2’),5’-bisphosphate nucleotidase, *CHB73_RS10980* a hypothetical protein and *CHB73_RS10975* an aldehyde ferredoxin oxidoreductase C-terminal domain-containing protein. C. Predicted components and topologies of the Hmc complex – the periplasmic 16-haem cytochrome *c* HmcA (light blue), C-tail containing iron-sulphur protein (grey), HybB-like quinol oxidase (cyan), small hydrophobic protein (dark grey), a NarI-related cytochrome *b* (bright blue) and a cytoplasmic iron-sulphur protein (grey). D. Genetic locus encoding the *hmc* gene cluster from *Desulfovibrio vulgaris* strain Hildenborough and *Solidesulfovibrio carbinolicus* strain DSM 3852. Genes encoding related proteins are shaded similarly, unrelated genes are shown in white. Genes *rrf1* and *rrf2* encode regulators of *hmc* expression (66). DVU_0538 is predicted to encode a family 2 AP endonuclease, *DVU_0528* a phosphatidylglycerophosphatase, *C3Y92_RS14895* a TOBE domain-containing protein, *C3Y92_RS14860* a universal stress protein, and *C3Y92_RS14850* a response regulator.

The second genomic locus was where a tail-anchored ferredoxin was encoded directly upstream of a quinol oxidase related to HybB and NrfD (Table S3, Fig 3). This tandem gene pair was found as part of a three, occasionally four gene module encoding in addition one or two periplasmic *c*-type cytochromes, frequently an octahaem *c*-type cytochrome from the tetrathionate reductase family (Table S3, Fig 3a,b). There was no pattern of conservation of genes flanking this cluster, and therefore it is unclear whether they participate as a module in a more extensive electron transport system. Blast analysis revealed a second occurrence of the tail-anchored iron-sulphur protein-quinol oxidase pair which was not found among examples in our dataset. Here the pairing is found as part of the high-molecular-weight cytochrome (Hmc) complex. This complex has been characterised from *Desulfovibrio vulgaris* subsp. *vulgaris* Hildenborough and comprises a unit of six redox proteins that contains, in addition to the high molecular weight cytochrome *c* (HmcA), tail-anchored iron-sulphur protein (HmcB) and quinol oxidase (HmcC), a hydrophobic peptide (HmcD), a NarI-related cytochrome *b* (HmcE) and a cytoplasmic GlpC superfamily iron-sulphur protein (HmcF) (Fig 3c,d) (41). Deletion analysis of the *hmc* operon results in impaired respiratory growth with hydrogen as sole electron donor, implicating it in electron transport between hydrogen and sulphate (42).

Interestingly, during our sequence analysis of the tail-anchored ferredoxin – quinol oxidase couple we found an unusual third instance of this pairing. However, in this case the iron-sulphur protein differed from those previously analysed because it lacks any predicted signal peptide, although the hydrophobic C-tail is conserved. Inspection of the flanking regions indicated that these two encoding genes are found immediately downstream of a gene coding for a periplasmic group A Fe-Fe hydrogenase. This gene locus was found in Gram-positive bacteria such as *Desulfotruncus alcoholivorax*, and Gram-negative *Proteus* species (Fig 4a,b). Sequence analysis indicates that the Fe-Fe hydrogenase has a twin arginine signal peptide that has a signal peptidase 2 cleavage site (Fig 4c) and is therefore likely to be lipid anchored. Since the tail-anchored iron-sulphur protein would be expected to have a similar topology to those that have both a Tat signal peptide and a C-tail, it is most likely co-exported with the signal peptide-bearing hydrogenase.

**Fig 4.**
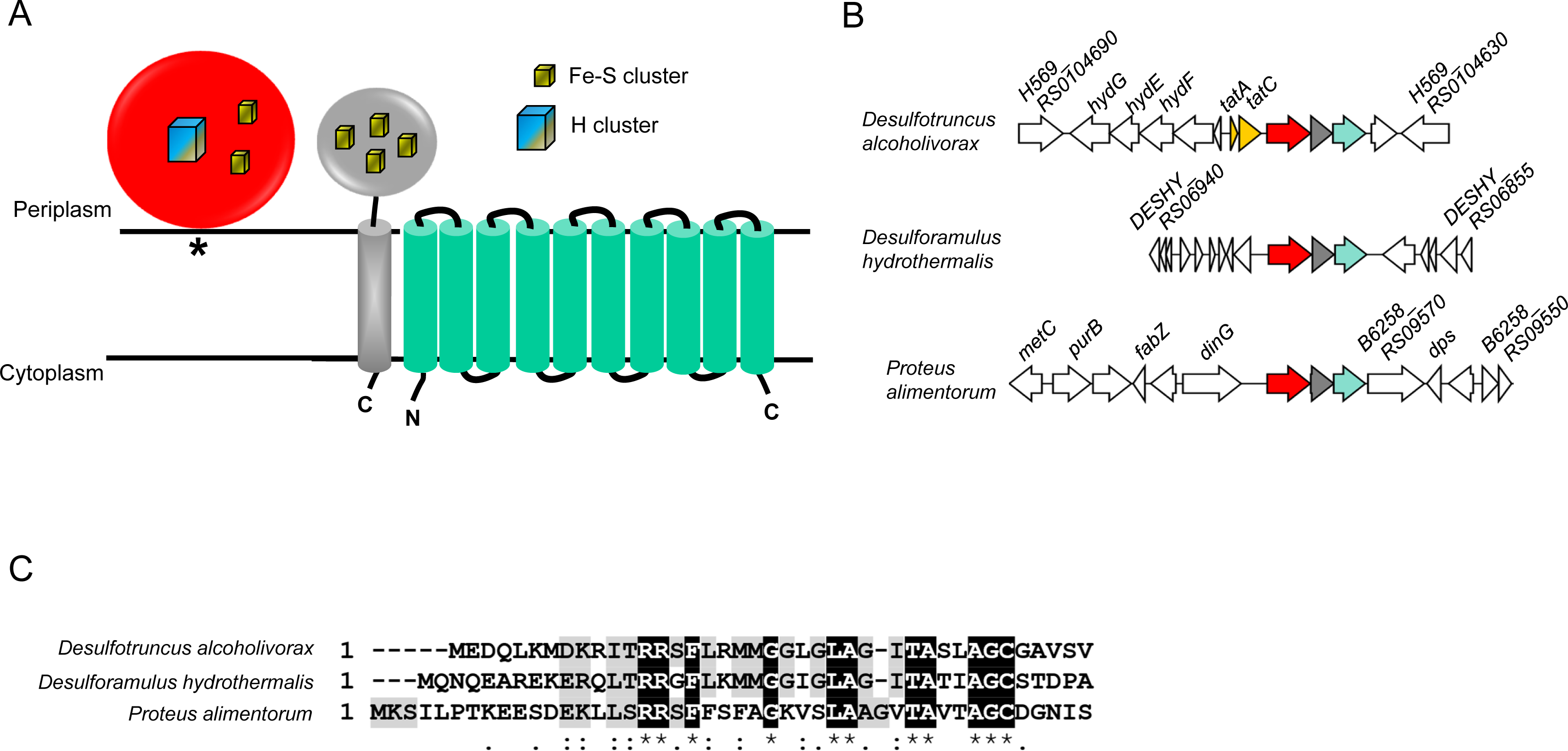
A C-tail iron-sulphur protein lacking a signal peptide may ‘hitchhike’ through the Tat pathway was a periplasmic Fe-Fe hydrogenase bearing a twin-arginine signal peptide. A. Predicted topologies of the three-protein module comprising a periplasmic group A Fe-Fe hydrogenase with a twin arginine lipoprotein signal peptide (red), a C-tail containing iron-sulphur protein (grey), and a HybB-like quinol oxidase (cyan). The asterisk indicates the lipid anchor. B. Example loci encoding the Fe-Fe hydrogenase, with genes coloured as in (A), and unrelated genes shown in white. For *Desulfotruncus alcoholivorax* strain DSM 16058, *H569_RS0104690* encodes a predicted HAMP domain-containing methyl-accepting chemotaxis protein, *hydG*, *hydE* and *hydF* encode Fe-Fe hydrogenase H-cluster maturation proteins, H569_RS19210 aspartate ammonia-lyase, H569_RS0104665 an iron-only hydrogenase system regulator, H569_RS0104635 a selenium metabolism-associated LysR family transcriptional regulator and H569_RS0104630 a cytoplasmic NADH-dependent Fe-Fe hydrogenase of group A6. For *Desulforamulus hydrothermalis* strain Lam5, genes *DESHY_RS06940*, *DESHY_RS06935*, *DESHY_RS06930*, *DESHY_RS06905*, *DESHY_RS06900*, *DESHY_RS06870*, *DESHY_RS06865* and *DESHY_RS06855* encode hypothetical proteins, *DESHY_RS06925* encodes a phage holin family protein, *DESHY_RS06920* an alpha/beta-type small acid-soluble spore protein, *DESHY_RS06915* a 4Fe-4S binding protein, *DESHY_RS06910* a YkvA family protein, *DESHY_RS06875* a cache domain-containing protein and *DESHY_RS06960* encodes SpoIIR. For *Proteus alimentorum* strain 08MAS0041, gene *metC* encodes cystathionine beta-lyase, *purB* adenylosuccinate lyase, *B6258_RS09605* a MmgE/PrpD family protein, *fabZ* 3-hydroxyacyl-ACP dehydratase, *B6258_RS09595* a LysR family transcriptional regulator, *dinG* an ATP-dependent DNA helicase, *B6258_RS09570* a TonB-dependent receptor, *dps* a DNA starvation/stationary phase protection protein, *B6258_RS09560* a RhtA threonine/homoserine exporter, *B6258_RS09555* a TIGR00730 family Rossman fold protein and *B6258_RS09550* a GNAT family N-acetyltransferase. C. Alignment of the twin arginine signal peptides of the Fe-Fe hydrogenases from the same three organisms.

### Proteins with other cofactors are also found in the dataset but are unlikely to be novel tail-anchored Tat substrates

In addition to iron-sulphur clusters, periplasmic proteins containing molybdopterin and cobalamin-based cofactors are also obligately Tat dependent (10, 43). The Tat pathway also exports substrates linked with other types of cofactor, including FAD, NAD/P, and copper ions, although some periplasmic proteins that bind these cofactors may be Sec dependent (43). We next searched dataset S1 for mention of any of these cofactors. From this we found five entries for FAD-containing proteins, of which one was a duplicate accession and was removed. We also found 43 entries for NAD-containing protein, of which ten were duplicates accessions and were removed. Finally, we found 21 entries for copper proteins, of which four were duplicate accessions. These proteins are listed in Table S4. None of the entries were annotated as molybdopterin or cobalamin containing.

Further analysis of the proteins in Table S4 revealed that none of the four FAD binding proteins present in dataset S1 were predicted to have any kind of signal peptide when reanalysed using SignalP 6.0. Moreover, the paired arginines in the N-terminal region were not conserved even among closely related sequences. The NAD binding proteins were only found among streptomycetes and further analysis indicated that the region annotated as the Tat signal peptide on the dataset output formed part of the nucleotide binding site, and again we conclude that they are cytoplasmically-located. Finally, the copper proteins identified in Table S4 have a CopC domain, predicted to bind one atom of copper per protein, and further analysis clearly predicts the presence of both an N-terminal signal peptide and a hydrophobic C-terminal stretch. However, SignalP-6.0 strongly predicts a Sec-targeting signal peptide and in agreement with this we noted that the RR motif was not conserved between otherwise closely related homologues. We therefore conclude that none of these cofactor-containing proteins are likely to be recognised by the Tat pathway.

### Bioinformatic sorting and analysis of other protein subsets

The extensive nature of dataset S1 precluded us from examining individual proteins, although we did take a closer inspection of some subsets. After hydrogenases, the largest number of proteins, 4,151, were annotated as hypothetical. To further classify these hypothetical proteins, we developed a script in Python allowing us to consider all of them as a “set”. From this only 1,852 unique hypothetical proteins were found, due to a large number of duplicated entries in the original dataset. From the 1,852 unique proteins, we ran the wanda.py script against RefSeq to obtain their descriptions. This expanded the list to 13,022 proteins because each RefSeq entry provides all the genomes that contain the requested WP. As each WP identifier groups all proteins that are 100% identical, we then used a custom script to select a representative assembly for each protein identifier, resulting in the final parsed output of 1,109 proteins that can be viewed in dataset S2.

From analysis of dataset S2 a further eight tail-anchored 4Fe-4S ferredoxins were found (Table S3), each of which was encoded at one of the genomic loci we observed above. In addition, WP_060849731.1 from *Methylobacterium aquaticum* was now annotated as a Rieske 2Fe-2S domain-containing protein. Analysing this protein by the structural prediction program Phyre2 (44), however, indicated that the hydrophobic region formed part of the Rieske fold rather than being predicted as a transmembrane helix. Moreover, amino acid substitutions were found in this hydrophobic region in closely related homologues that reduced the hydrophobicity while not altering the structural prediction. We therefore conclude that this is unlikely to be a tail-anchored Tat substrate protein.

Two additional FAD/NAD(P)-binding oxidoreductase proteins were present in the reannotated hypothetical list in dataset S2. One of these, WP_020186848.1, lacks a predicted signal peptide using SignalP 6.0. By contrast, BLAST analysis of WP_052700213.1 from *Methylocucumis oryzae* indicates that the twin-arginine signal peptide is conserved among close homologues. However, all of the close homologues are substantially longer than WP_052700213.1, with no predicted C-terminal hydrophobic stretches. Closer inspection reveals a frameshift in WP_052700213.1 giving rise to a string of leucines and isoleucines near the truncated C-terminus of this protein accession – it is not clear at present whether this represents a genuine frameshift or a sequencing error. Finally, a further 26 CopC homologues were identified in dataset S2, but again these were predicted to be Sec substrates using SignalP-6.0.

After hypothetical proteins, ‘inhibition of morphological differentiation proteins’ and ‘morphological differentiation-associated protein’ was one of the most frequent annotations, collectively appeared in 813 entries in dataset S1. Analysis showed that these are all proteins are of the haloacid dehalogenase (HAD)-like hydrolase family. A further example was directly annotated as HAD-superfamily hydrolase in dataset S1, with another six HAD annotations also present in dataset S2. When we analysed these further, we noted that the C-terminal hydrophobic stretch was conserved, but that the proteins are predicted to lack signal peptides when analysed by SignalP 6.0. Instead, the paired arginines form part of the HAD signature motif I, accounting for their conservation.

Ten accessions in the dataset S1 had ‘Tat pathway protein signal’ in their annotation (one of which was a duplicate) and 49 contained ‘twin-arginine translocation’, of which 14 were duplicate entries. Three additional proteins with ‘‘twin-arginine translocation’ were also present in dataset S2. These proteins are listed in Table S5. Seven of the proteins with ‘Tat pathway protein signal’ were alginate lyase family proteins. Closely related proteins to these entries also had predicted C-tails but the twin arginines in the signal sequence were not conserved and are therefore unlikely to be Tat substrates. Almost all of the entries under ‘twin-arginine translocation’ were from *Stenotrophomonas* and are predicted to be subunits of gluconate-2-dehydrogenase. While the twin arginine signal peptide appears to be quite widely conserved across this protein family, the hydrophobic tail region forms part of the gluconate_2-dh3 fold and we identified substitutions in this region in homologous proteins that abolished the prediction of a TMH. We conclude that these entries are unlikely to be tail-anchored Tat substrates.

### Proteases and peptidases are found among the datasets

The annotations ‘peptidase’ and ‘protease’ were commonly found in dataset S1. Further analysis indicated that these fall into several different groups. Mycosin protease was one of the most frequently identified. This is a serine protease associated with the type VII secretion system and there were 116 entries for ‘mycosin’ and 186 for ‘type VII’ among the annotation in dataset S1. Some of the entries in datasets S1 and S2 were annotated as ‘S8 peptidase’, and on further analysis these were also mycosin protease homologues. Mycosin is a known extracellular tail-anchored protein that is synthesised with an N-terminal signal peptide (45, 46). There are now over 60,000 mycosin protein sequences available in NCBI and examination of 100 sequences at random revealed that although they were all predicted to have N-terminal signal peptides, almost all of them lack paired arginines in the n-region. Mycosins are therefore unlikely to engage with the Tat pathway.

‘S1 peptidase’ was also found among the annotation in dataset S1. S1 peptidases are a large family of serine endopeptidases related to chymotrypsin A (47). We found 19 accessions with this annotation in dataset S1 (of which three were duplicates) and a further three in dataset S2 - these are listed in Table S6). We also noted that several dataset S1 accessions were annotated as ‘serine protease’ and further analysis of these revealed an additional 25 S1 peptidases, of which one was a duplicate (Table S6). Strikingly, almost all of the S1 peptidases were from *Streptomyces*. This genus of bacteria are prolific secretors of proteins and make extensive use of the Tat pathway (48–51). Many *Streptomyces* Tat substrates lack cofactors and their homologues in other bacteria are Sec substrates (48, 49), suggesting that these S1 peptidases are plausible candidates for Tat-dependent C-tail proteins. A sequence alignment of seven *Streptomyces* S1 peptidases in Fig S1A highlights the conservation of the signal peptide and C-tail regions, and an Alphafold structure prediction in Fig S1B clearly suggests that the C-tail is a distinct hydrophobic helix rather than forming an integral part of the folded catalytic domain.

### *Streptomyces* D-alanyl-D-alanine carboxypeptidases are plausible Tat-dependent tail-anchored protein candidates

The annotation D-alanyl-D-alanine carboxypeptidase was found in 115 instances in dataset S1 (of which 34 were duplicate accessions), and a further two were found in dataset S2. These proteins are listed in Table S7. D-alanyl-D-alanine carboxypeptidases remove the terminal d-alanine residue from pentapeptide side chains of peptidoglycan, influencing the level of peptidoglycan crosslinking (52). Most of the entries in the datasets are from the *Streptomyces* genus, and further analysis indicated that they are all ‘DacC’ type enzymes. The DacC protein from the model streptomycete *S. coelicolor* has been identified experimentally as part of the membrane proteome and is predicted to be a Tat substrate (53). In this context it has been noted that the cell walls of *Streptomyces tat* mutants are particularly fragile, consistent with a defect in peptidoglycan synthesis and/or remodelling (48, 49, 51). Surprisingly, *S. coelicolor* DacC (CAB89066.1) was not identified by our bioinformatic search, but inspection of the sequence reveals that it fulfils the criteria for a Tat-dependent tail-anchored protein and Fig S2 shows that it shares significant homology with other d-alanyl-d-alanine carboxypeptidases from dataset S1.

### Sortase substrates are found among the accessions in the datasets

Sortases are transpeptidases found in Gram-positive bacteria that fall into six classes, A-F (54, 55). They covalently attach proteins to the peptide sidechains of peptidoglycan (54). Sortase substrates have a C-terminal transmembrane helix that is preceded by a sorting motif, often LPXTG (56). Sortase cleaves between the threonine and glycine, ultimately resulting in the attachment of the threonine to the free amino group of the lipid II-bound cell wall precursor. Most sortase substrates are exported across the membrane by the Sec pathway, and to our knowledge no naturally-occurring Tat-dependent sortase substrate has been described. However it has been reported that fusing a Tat signal peptide to a Sec-dependent sortase substrate re-routes it to the Tat pathway but does not affect its cell wall anchoring (57).

When we further analysed the accessions that were annotated as ‘peptidase’ in dataset S1, we noted that a small subset of seven proteins, all from *Streptomyces*, had a predicted sortase recognition motif which was not part of their original annotation (Table S8; shown in the alignment in Fig S3). SignalP-6.0 strongly predicted that the presence of twin arginine signal peptides suggesting that these proteins may engage with the Tat pathway during their biogenesis.

We next turned our attention to other predicted sortase substrates. Fourteen proteins in dataset S1 were noted to have an ‘LPXTG’ annotation of which two were duplicates, these are listed in Table S8. A further 15 proteins with ‘LPXTG’ annotation (Table S8) were also identified in dataset S2. *Streptomyces* genomes usually encode at least two sortase enzymes, minimally one from class E and one from class F (58). SrtE recognises and cleaves at a LAETG sorting motif (59). We identified 19 proteins with ‘LAETG’ annotation in dataset S2 and these are also listed in Table S8.

Analysis of the entries in Table S8 indicates that 12 of the ‘LAETG’ annotated proteins were from the SCO1860 family (Fig S4). When we inspected the sequence of the founder member of this family, SCO1860 from *S. coelicolor* (accession QFI42043.1) it was not recognised to have a Tat signal peptide most likely because the predicted sequence has a very long n-region prior to the twin arginine motif. A more recent genome sequence of *S. coelicolor* from 2021 has assigned a TTG start codon and a shorter n-region (WP_038535008.1) that now meets the criteria for inclusion as a candidate Tat-dependent protein synthesised with a C-terminal tail. It therefore appears that these features are well conserved across the protein family.

### Assessing the presence of Tat signal peptides and C-tails using protein fusions

The bioinformatic analysis above has identified several candidate Tat-dependent C-tail proteins, including peptidases, carboxypeptidases and sortase substrates. To experimentally confirm these predictions, we constructed reporter fusions for analysis in *E. coli*. The *E. coli* cell wall amidases, AmiA and AmiC are native Tat substrates, and their mislocalisation in *tat* mutant strains results in a defect in cell wall remodelling resulting in sensitivity to killing by detergents such as SDS (34, 60). A similar phenotype is also observed if the *amiA* and *amiC* genes are deleted in a *tat*^+^ background, which can be rescued by production of AmiA *in trans* (60, 61). This forms the basis of a sensitive screen for Tat signal peptides – the native signal sequence of AmiA can be replaced by other targeting sequences and if they engage with the Tat pathway they permit the *amiA*/*amiC* mutant strain to grow in the presence of SDS (18). It should be noted that AmiA can also be exported in a functional form by the Sec pathway (18, 60). By repeating the same growth assays in *amiA*/*amiC* mutant strain that is additionally deleted for *tat* genes this will reveal whether growth on SDS arises from engagement of the fusion with the Sec pathway rather than being strictly Tat dependent.

To this end, we fused the signal peptide coding regions of 16 protein candidates to the mature sequence of AmiA and assessed their ability to mediate Tat-dependent export. The candidates are listed in Table 2 and include an S1 family peptidase (candidate 1); two predicted sortase-anchored peptidases (candidates 4 and 6); six additional predicted sortase substrates (candidates 7, 10, 11, 12, 13 and 14 including two - candidates 13 and 14 - from the SCO1860 family) and two carboxypeptidases (candidates 15 and 16). We also selected a further five candidates from the dataset S1 – two HtaA-domain proteins (candidates 3 and 5), a YcnI family protein (candidate 2), a terpene cyclase/mutase family protein (candidate 8) and a DUF4349 domain protein (candidate 9). In these experiments the signal peptide of the *E. coli* Tat substrate TorA is used as a positive control, as fusions to this signal peptide are strictly Tat dependent due to the presence of two positive charges in the c-region that act as a Sec-avoidance motif (17).

**Table 2.**
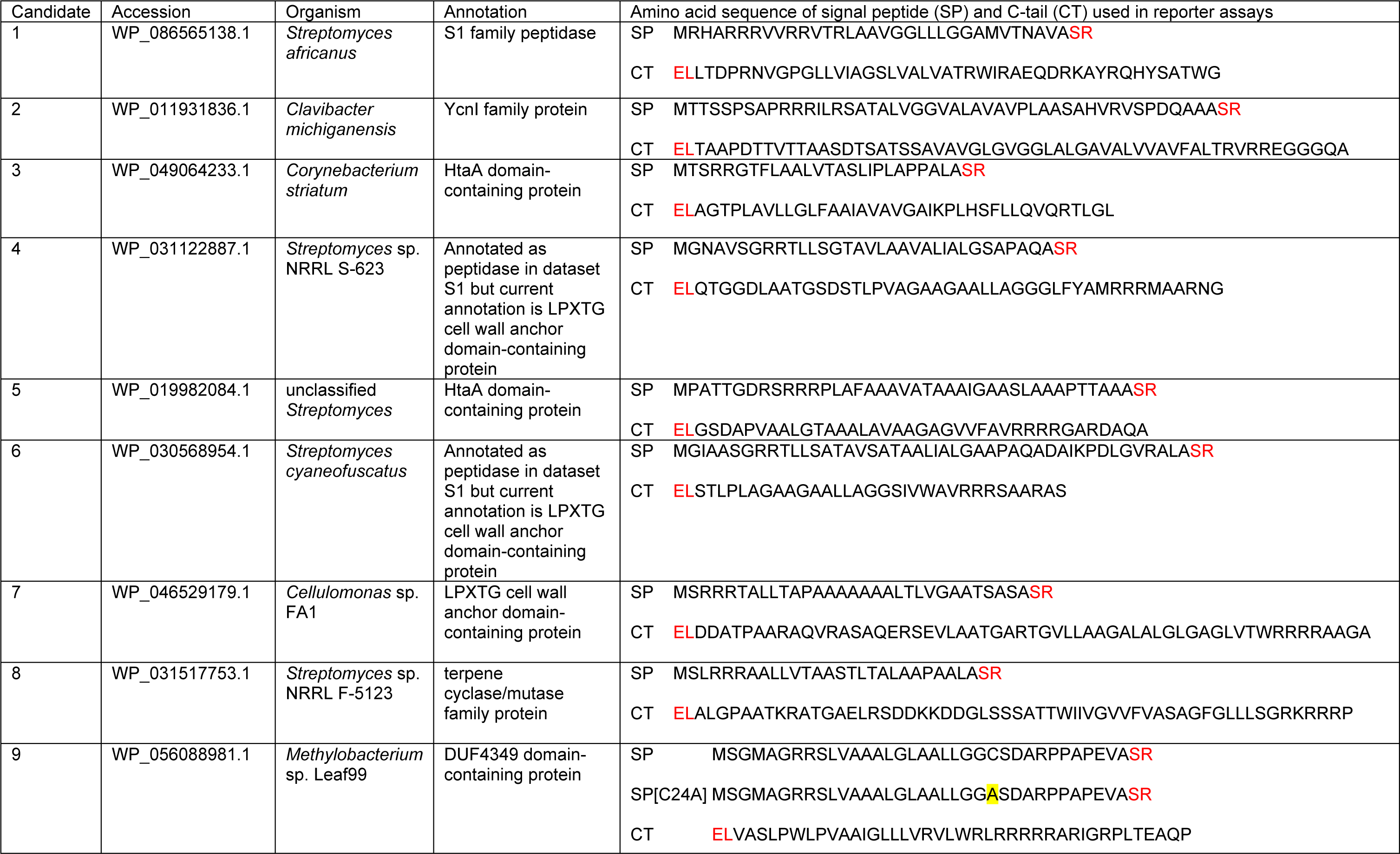
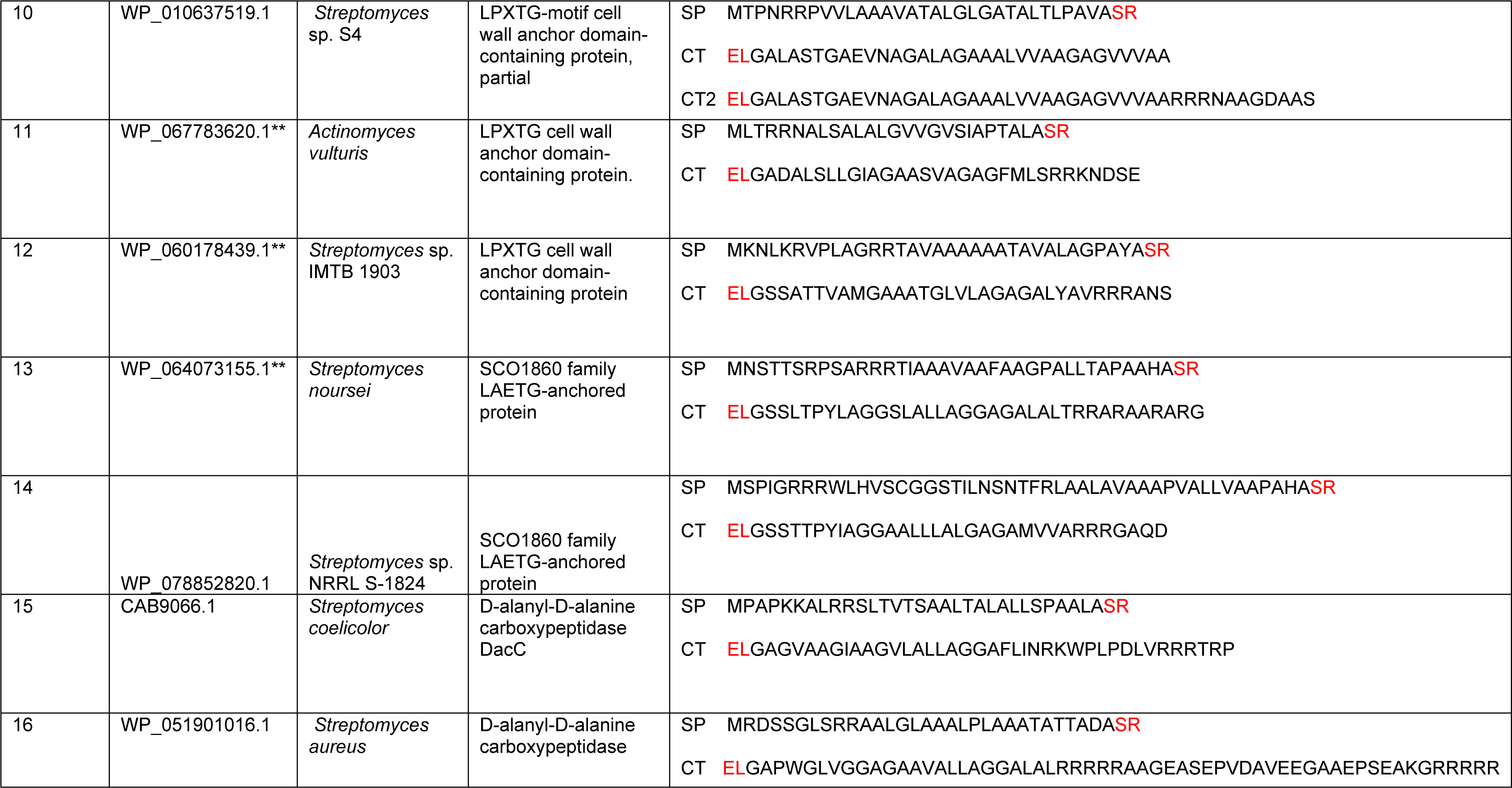
Candidate Tat-dependent tail-anchored proteins identified in this study that were selected for experimental validation using AmiA and SufI reporter assays. The amino acid residues encoded by the restriction site at the fusion junctions are shown in red. The residue highlighted in yellow for candidate 9 indicates substitution of the cysteine residue in the lipobox for alanine.

As shown in Fig 5A,B the TorA signal peptide fused to AmiA supports growth of the *amiA*/*amiC* mutant on SDS when Tat is functional but not when the pathway is inactivated. Strains producing the signal peptides of candidates 1, 2, 10, 11, 12, 13 and 15 fused to AmiA showed a growth pattern similar to those producing TorA-AmiA, i.e. robust growth of the *E. coli amiA*/*amiC* mutant strain in the presence of SDS but little or no growth for the *amiA*/*amiC*/*tat* mutant strain (Fig 5A). By contrast, the signal peptides of candidates 3, 5, 7, 8, 9, 14 and 16 were unable to mediate any detectable export of AmiA as none of the fusions were able to restore growth of the *amiA*/*amiC* mutant in SDS-containing media (Fig S5).

**Fig 5.**
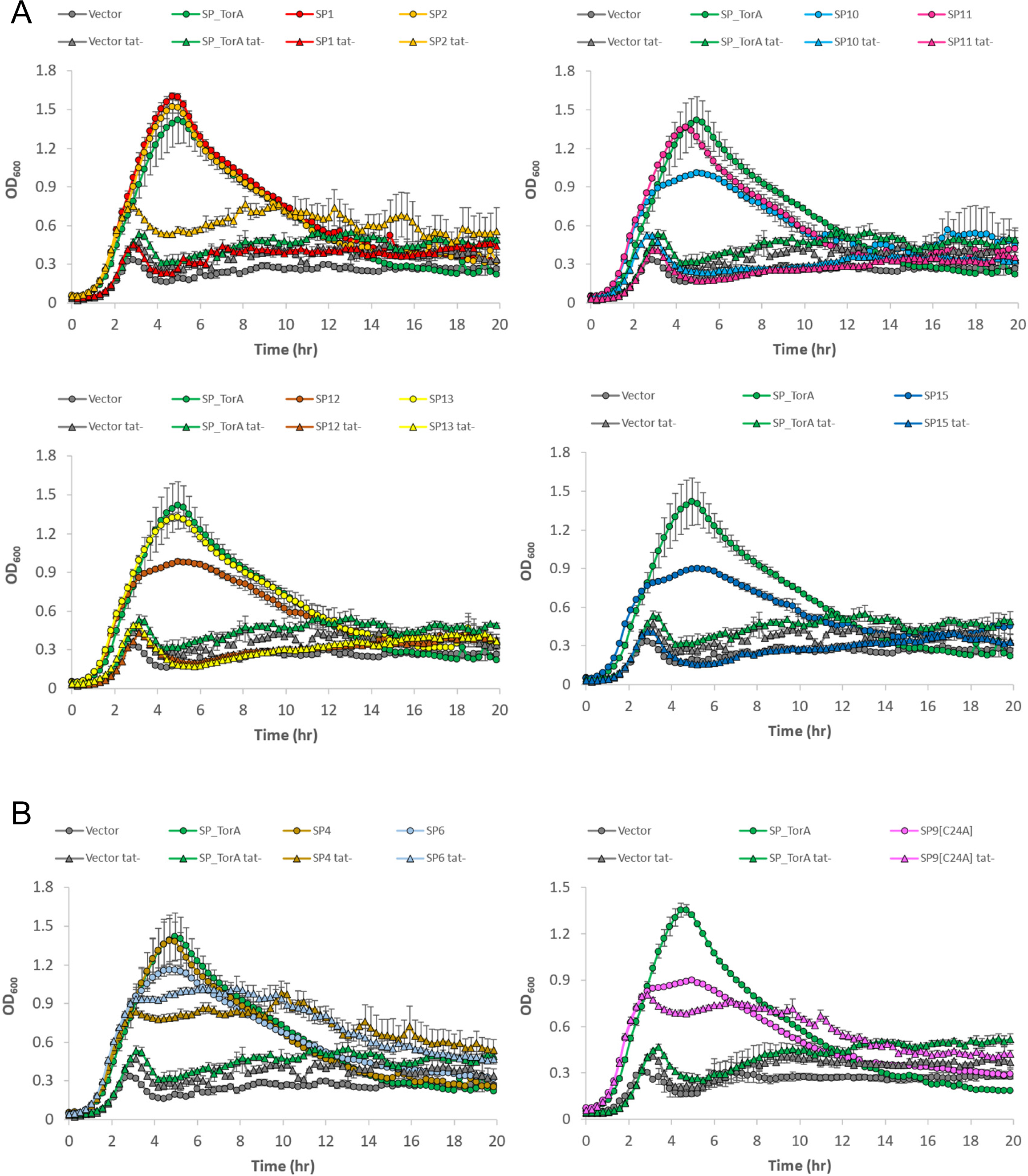
The signal peptides of candidates 1, 2, 10, 11, 12, 13 and 15 mediate Tat-dependent export of AmiA. Plasmids encoding the indicated signal peptide (SP) - AmiA fusions were introduced into *E. coli* strains MC4100 Δ*amiA* Δ*amiC* (circles) and MC4100 Δ*amiA* Δ*amiC* Δ*tatABC* (triangles and labelled tat-) and grown for 20 h in LB in the presence of 0.5% SDS without shaking in a plate reader. Growth curves correspond to an averaged triplicate set ± one SD (standard deviation). For clarity, growth curves are displayed either in groups of 2 or individually alongside the positive (“SP_TorA”) and negative (“Vector”, i.e., pUniAmiA) controls in each plot. A. Growth curves of strains expressing SP-AmiA fusions that are fully dependent on Tat for export. B. Growth curves of strains expressing SP-AmiA fusions that are exported in both *tat*^+^ and *tat*^-^ strains. Growth curves of strains expressing non-exported SP-AmiA fusions are shown in Fig. S5.

On closer inspection we noted that the signal peptide of candidate 9, the DUF4349 domain protein was predicted to harbour a lipobox sequence. While lipoproteins are compatible with export by the Tat pathway (62), the amidase reporter assay does not work well for membrane-anchored Tat substrates, presumably because tethering to the membrane prevents AmiA from interacting with the peptidoglycan substrate (7). We therefore mutated the lipobox cysteine of this signal peptide to alanine. Fig 5B indicates that the C24A variant of this signal peptide could now support growth of the *amiA*/*amiC* strain when fused to AmiA. However, a similar level of growth was observed in an *amiA*/*amiC*/*tat* background, indicating significant engagement of this construct with the Sec pathway. A similar pattern of growth behaviour was also observed with the signal peptides of candidates 4 and 6 which were also able to mediate export of AmiA in the absence of a functional Tat system (Fig 5B). While it remains possible that these signal peptides are Tat dependent in their native organisms, we are unable to confirm their engagement with the Tat pathway using this heterologous system.

To explore the presence of C-tails on these candidate proteins we used the *E. coli* Tat substrate SufI as a reporter. Native SufI is a soluble periplasmic protein but it can be anchored to the periplasmic side of the inner membrane if a transmembrane segment is fused to its C-terminus (5). We therefore fused the predicted C-tail regions of each protein to SufI and determined the subcellular location of each of the SufI fusion proteins. Fig 6 shows that very little native SufI can be detected in urea-washed membranes, but that if it is fused to the C-tail region of FdnH it is primarily found in the washed membrane fraction, as expected. The C-tail regions of all protein candidates, except for 10 and 11 were similarly able to mediate urea-resistant integration of SufI into the membrane (Fig 6A, B). In several cases some residual SufI was also detected in the soluble fraction, but this was generally as smaller forms than the integrated fusion and probably arises from proteolytic processing of these non-native constructs at the C-tail region.

**Fig 6.**
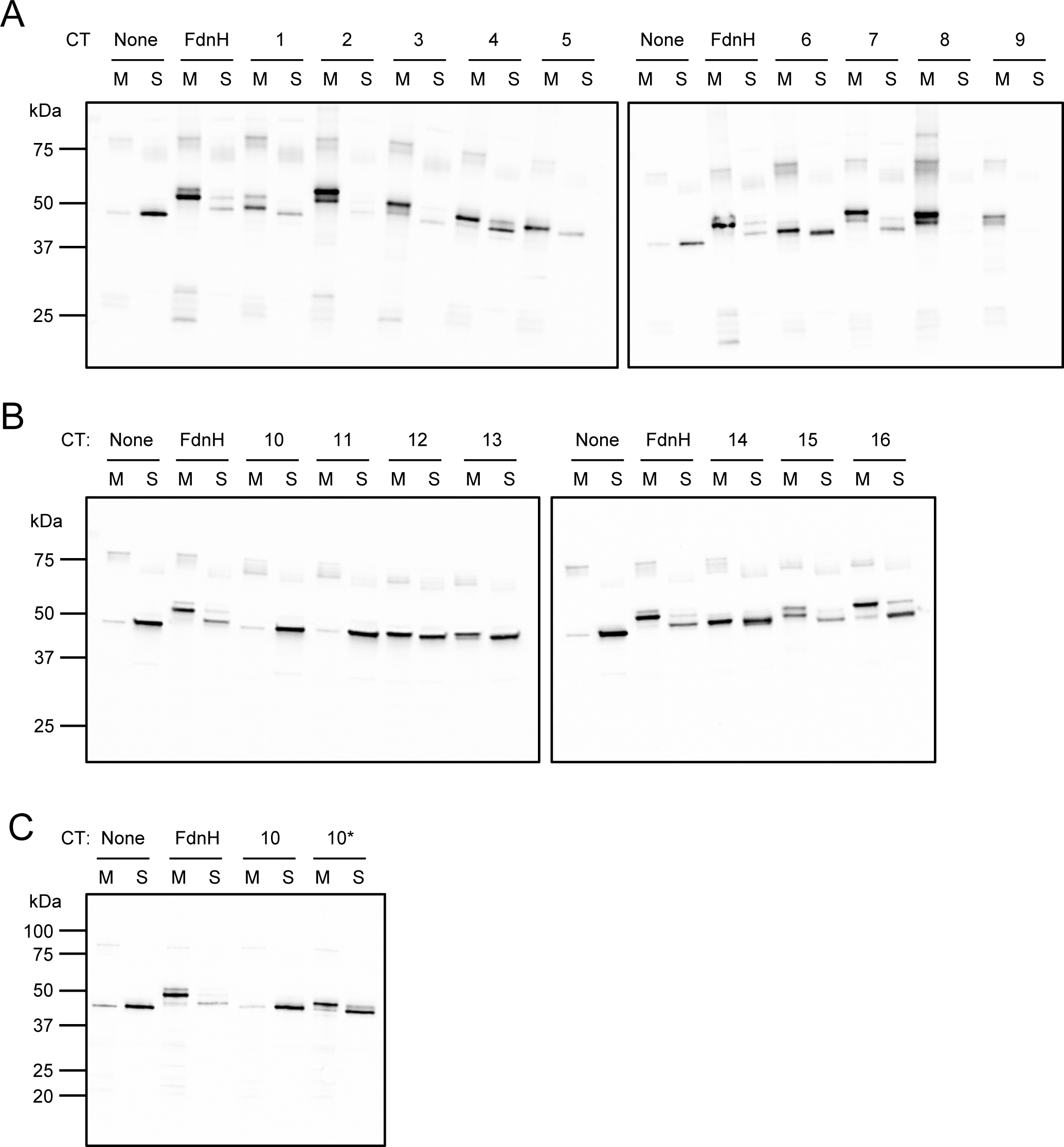
Subcellular localisation of SufI C-tail fusions expressed in an *E. coli* Δ*sufI* strain. Cultures of *E. coli* strain NRS-3 (Δ*sufI*) expressing the indicated SufI/C-tail (“CT”) fusions were fractionated into total soluble (cytoplasm + periplasm, “S”) and urea-washed membrane (“M”) fractions, which were then analysed by western blotting using specific anti-SufI antibodies. “None”: WT SufI with no fused C-tail; “FdnH”: SufI fused with the C-tail of FdnH (positive control for membrane insertion); “1-16”: C-tails from the protein targets listed in Table 2; “10*”: amended, extended C-tail for protein target 10 (Table 2). Due to the number of samples and the consequent use of multiple blots, each blot includes an independent set of controls for membrane insertion.

When we re-examined the sequence of candidate 10 we noted that this accession (WP_010637519.1) was now annotated as obsolete. Blast analysis revealed that the closest homologue to this accession is WP_129815067.1, a protein with two predicted HtaA domains from *Streptomyces albidoflavus*. The two proteins share 99.8% identity across the 503 amino acids of WP_010637519.1, differing by a single amino acid at position 442. However, WP_129815067.1 and other homologues are all 514 amino acids in length, with additional sequence following the predicted C-tail region. We therefore extended the sequence of the C-tail encoded by the candidate 10 fusion to include this additional sequence and repeated the fractionation. Fig 6C indicates that this extended sequence is now able to direct membrane integration of SufI and we conclude that this is also a *bona fide* transmembrane segment.

Taken together the results of the fusion protein analysis have confirmed six novel Tat-dependent C-tail proteins - WP_086565138.1, an S1 family peptidase from *Streptomyces africanus*, WP_011931836.1, a YcnI family protein from *Clavibacter michiganensis*, WP_129815067.1, a HtaA domain LPXTG protein from *Streptomyces albidoflavus*, WP_060178439.1, LPXTG cell wall anchor domain-containing protein from *Streptomyces* sp. IMTB 1903, WP_064073155.1, a SCO1860 family LAETG-anchored protein from *Streptomyces noursei* and CAB9066.1, the D-alanyl-D-alanine carboxypeptidase DacC from *Streptomyces coelicolor*.

## DISCUSSION

Here we have taken a bioinformatic approach to identify novel C-tail anchored proteins that are integrated into the membrane by the twin-arginine translocase. Prior work had revealed the presence of five such proteins in *E. coli*, including the iron-sulphur cluster containing small subunits of periplasmic Ni-Fe hydrogenases and formate dehydrogenases, and the electron transfer protein HybA (5). Searching for proteins with a predicted N-terminal Tat signal peptide and a hydrophobic stretch in the last 50 amino acids identified over 34,000 proteins that met these criteria.

Analysis of the output revealed that many of the proteins were either small subunits of periplasmic hydrogenases or homologues of HybA, validating our approach. In addition to these known iron-sulphur proteins we also identified related proteins that were not associated with hydrogenase gene clusters. We found a pairing with a *b*-type cytochrome that form an apparent stand-alone electron transport module, and a second pairing with a quinol oxidase that were usually encoded next to one or more *c*-type cytochromes. Further analysis with these novel tail-anchored ferredoxins identified that they can also form part of a larger electron transport cluster such as the high-molecular-weight cytochrome complex. This analysis also revealed an example of a tail-anchored ferredoxin lacking any signal peptide that is likely exported as a partner with a Tat-dependent Fe-Fe hydrogenase.

Our output additionally identified many proteins that were not predicted to contain any cofactors. Manually sorting these into groups indicated that many candidate proteins were from the *Streptomyces* genus. This is perhaps not surprising given that these organisms have the largest number of Tat substrates identified to date (48, 49, 63). The high percentage of chromosomal G+C also skews a bias towards arginine codons over lysine codons, the latter of which comprise only A+T bases.

We noted that proteases, carboxypeptidases and sortase substrates were enriched among the output proteins. To assess Tat dependence of a selection of these proteins, we determined the ability of the predicted signal peptides from 16 candidates to mediate Tat-dependent export of an *E. coli* reporter protein, finding that at least seven of these could functionally engage with the Tat pathway. Of these seven, six also had a C-terminal amino acid stretch that could serve as an integral membrane anchor, and we conclude based on these findings that they represent novel Tat-dependent C-tail proteins. Among the proteins we have verified include four predicted sortase substrates. Sortase substrates are transiently membrane-anchored during their biogenesis, prior to cleavage at a motif preceding the anchor sequence and covalent attachment of the globular domain to peptidoglycan (56). Our findings indicate that Tat dependent sortase substrates may be abundant in actinobacteria.

One of the limitations of this study is that it does not directly identify Tat substrate proteins that have a C-tail but lack a signal peptide because they are exported with Tat-dependent partner proteins. Such candidates would be difficult to identify bioinformatically as it would require a more complex search where a predicted Tat substrate was encoded in the genetic neighbourhood of a tail anchored protein. In this context, a previous search for candidate tail-anchored proteins encoded by *Streptomyces coelicolor* identified 20 such proteins that lacked any identifiable signal peptide and could potentially be exported through binding to a Tat targeting partner protein. However only six of them were predicted to have an N-out configuration (64) and from our analysis none of these are encoded next to candidate Tat substrates.

In conclusion, six new tail-anchored membrane proteins that are integrated by the Tat pathway have been validated experimentally. None of these proteins are predicted to bind redox cofactors and they represent the first such tail-anchored proteins that are not involved in electron transport. We anticipate that the approach taken here could be used to find further tail anchored Tat substrates.

## FUNDING INFORMATION

This work was supported by UKRI Biotechnology and Biological Sciences Research council Grant/Award Number BB/S005307/1 and Medical Research Council Grant/Award Number MR/S009213/1.

## ACKNOWLEDGEMENTS

We thank Dr Stephen Garrett for his help with generating the Clinker outputs, Dr Giuseppina Mariano and Dr Jorge Camarero-Vera and for bioinformatics and database help and Profs Phillip Stansfeld, Ben Berks, Frank Sargent and Dr Jon Cherry for helpful discussion.

## AUTHOR CONTRIBUTIONS

TP conceived the project. GC, JJG, and ES conducted experiments and JJG, ES and TP analysed data. TP wrote the manuscript with input from ES. All authors approved the submission.

## CONFLICTS OF INTEREST

The authors declare no conflict of interests

## Notes

### Competing Interest Statement

The authors have declared no competing interest.

## References

1. Wallin E, von Heijne G. Genome-wide analysis of integral membrane proteins from eubacterial, archaean, and eukaryotic organisms. Protein Sci 1998;7(4):1029–1038.

2. Krogh A, Larsson B, von Heijne G, Sonnhammer EL. Predicting transmembrane protein topology with a hidden Markov model: application to complete genomes. J Mol Biol 2001;305(3):567–580.

3. Kuhn A, Koch HG, Dalbey RE. Targeting and Insertion of Membrane Proteins. EcoSal Plus 2017;7(2).

4. De Geyter J, Smets D, Karamanou S, Economou A. membrane translocases and insertases. Subcell Biochem 2019;92:337–366.

5. Hatzixanthis K, Palmer T, Sargent F. A subset of bacterial inner membrane proteins integrated by the twin-arginine translocase. Mol Microbiol 2003;49(5):1377–1390.

6. Bachmann J, Bauer B, Zwicker K, Ludwig B, Anderka O. The Rieske protein from *Paracoccus denitrificans* is inserted into the cytoplasmic membrane by the twin-arginine translocase. FEBS J 2006;273(21):4817–4830.

7. Tooke FJ, Babot M, Chandra G, Buchanan G, Palmer T. A unifying mechanism for the biogenesis of membrane proteins co-operatively integrated by the Sec and Tat pathways. Elife 2017;6:e26577.

8. Passmore IJ, Dow JM, Coll F, Cuccui J, Palmer T, Wren BW. The ferric citrate regulator, FecR, is translocated across the bacterial inner membrane via a unique Twin-arginine transport dependent mechanism. J Bacteriol 2020;202:e00541–19.

9. DeLisa MP, Tullman D, Georgiou G. Folding quality control in the export of proteins by the bacterial twin-arginine translocation pathway. Proc Natl Acad Sci U S A 2003;100(10):6115–6120.

10. Berks BC. A common export pathway for proteins binding complex redox cofactors? Mol Microbiol 1996;22(3):393–404.

11. Dilks K, Rose RW, Hartmann E, Pohlschroder M. Prokaryotic utilization of the twin-arginine translocation pathway: a genomic survey. J Bacteriol 2003;185(4):1478–1483.

12. Bendtsen JD, Nielsen H, Widdick D, Palmer T, Brunak S. Prediction of twin-arginine signal peptides. BMC Bioinformatics 2005;6:167.

13. Stanley NR, Palmer T, Berks BC. The twin arginine consensus motif of Tat signal peptides is involved in Sec-independent protein targeting in *Escherichia coli*. J Biol Chem 2000;275(16):11591–11596.

14. Rollauer SE, Tarry MJ, Graham JE, Jaaskelainen M, Jager F, et al. Structure of the TatC core of the twin-arginine protein transport system. Nature 2012;492(7428):210-214.

15. Yahr TL, Wickner WT. Functional reconstitution of bacterial Tat translocation *in vitro*. EMBO J 2001;20(10):2472–2479.

16. Tsirigotaki A, De Geyter J, Sostaric N, Economou A, Karamanou S. Protein export through the bacterial Sec pathway. Nat Rev Microbiol 2017;15(1):21–36.

17. Cristobal S, de Gier JW, Nielsen H, von Heijne G. Competition between Sec- and TAT-dependent protein translocation in *Escherichia coli*. EMBO J 1999;18(11):2982–2990.

18. Huang Q, Palmer T. Signal peptide hydrophobicity modulates interaction with the twin-arginine translocase. mBio 2017;8(4):e00909–17.

19. Bogsch E, Brink S, Robinson C. Pathway specificity for a delta pH-dependent precursor thylakoid lumen protein is governed by a ‘Sec-avoidance’ motif in the transfer peptide and a ‘Sec-incompatible’ mature protein. EMBO J 1997;16(13):3851–3859.

20. Tullman-Ercek D, DeLisa MP, Kawarasaki Y, Iranpour P, Ribnicky B, et al. Export pathway selectivity of *Escherichia coli* twin arginine translocation signal peptides. J Biol Chem 2007;282(11):8309–8316.

21. Berks BC, Sargent F, Palmer T. The Tat protein export pathway. Mol Microbiol 2000;35(2):260–274.

22. Palmer T, Sargent F, Berks BC. The Tat Protein Export Pathway. EcoSal Plus 2010;4(1).

23. Rodrigue A, Chanal A, Beck K, Muller M, Wu LF. Co-translocation of a periplasmic enzyme complex by a hitchhiker mechanism through the bacterial Tat pathway. J Biol Chem 1999;274(19):13223–13228.

24. Stanley NR, Sargent F, Buchanan G, Shi J, Stewart V, et al. Behaviour of topological marker proteins targeted to the Tat protein transport pathway. Mol Microbiol 2002;43(4):1005–1021.

25. Sargent F, Ballantine SP, Rugman PA, Palmer T, Boxer DH. Reassignment of the gene encoding the *Escherichia coli* hydrogenase 2 small subunit--identification of a soluble precursor of the small subunit in a *hypB* mutant. Eur J Biochem 1998;255(3):746–754.

26. Jormakka M, Tornroth S, Byrne B, Iwata S. Molecular basis of proton motive force generation: structure of formate dehydrogenase-N. Science 2002;295(5561):1863-1868.

27. Beaton SE, Evans RM, Finney AJ, Lamont CM, Armstrong FA, et al. The structure of hydrogenase-2 from *Escherichia coli*: implications for H(2)-driven proton pumping. Biochem J 2018;475(7):1353–1370.

28. Rose RW, Bruser T, Kissinger JC, Pohlschroder M. Adaptation of protein secretion to extremely high-salt conditions by extensive use of the twin-arginine translocation pathway. Mol Microbiol 2002;45(4):943–950.

29. Saha CK, Sanches Pires R, Brolin H, Delannoy M, Atkinson GC. FlaGs and webFlaGs: discovering novel biology through the analysis of gene neighbourhood conservation. Bioinformatics 2021;37(9):1312–1314.

30. Gilchrist CLM, Chooi YH. Clinker & clustermap.js: Automatic generation of gene cluster comparison figures. Bioinformatics 2021;btab007. doi: 10.1093/bioinformatics/btab007.

31. Teufel F, Almagro Armenteros JJ, Johansen AR, Gislason MH, Pihl SI, et al. SignalP 6.0 predicts all five types of signal peptides using protein language models. Nat Biotechnol 2022;40(7):1023–1025.

32. Hallgren J, Tsirigos KD, Pedersen MD, Almagro Armenteros JJ, Marcatili P, et al. DeepTMHMM predicts alpha and beta transmembrane proteins using deep neural networks. BioRxiv 2022;10.1101/2022.04.08.487609.

33. Casadaban MJ, Cohen SN. Lactose genes fused to exogenous promoters in one step using a Mu-*lac* bacteriophage: *in vivo* probe for transcriptional control sequences. Proc Natl Acad Sci U S A 1979;76(9):4530–4533.

34. Stanley NR, Findlay K, Berks BC, Palmer T. *Escherichia coli* strains blocked in Tat-dependent protein export exhibit pleiotropic defects in the cell envelope. J Bacteriol 2001;183(1):139–144.

35. Huang Q, Alcock F, Kneuper H, Deme JC, Rollauer SE, et al. A signal sequence suppressor mutant that stabilizes an assembled state of the twin arginine translocase. Proc Natl Acad Sci U S A 2017;114:E1958–E1967.

36. Ize B, Coulthurst SJ, Hatzixanthis K, Caldelari I, Buchanan G, et al. Remnant signal peptides on non-exported enzymes: implications for the evolution of prokaryotic respiratory chains. Microbiology 2009;155(Pt 12):3992–4004.

37. Jack RL, Buchanan G, Dubini A, Hatzixanthis K, Palmer T, Sargent F. Coordinating assembly and export of complex bacterial proteins. EMBO J 2004;23(20):3962–3972.

38. Liu H, Naismith JH. An efficient one-step site-directed deletion, insertion, single and multiple-site plasmid mutagenesis protocol. BMC Biotechnol 2008;8:91.

39. Severi E, Bunoro Batista M, Lannoy A, Stansfeld PJ, Palmer T. Characterization of a TatA/TatB binding site on the TatC component of the *Escherichia coli* twin arginine translocase. Microbiology 2023;169(2):doi: 10.1099/mic.0.001298.

40. Buchanan G, de Leeuw E, Stanley NR, Wexler M, Berks BC, et al. Functional complexity of the twin-arginine translocase TatC component revealed by site-directed mutagenesis. Mol Microbiol 2002;43(6):1457–1470.

41. Rossi M, Pollock WB, Reij MW, Keon RG, Fu R, Voordouw G. The *hmc* operon of *Desulfovibrio vulgaris* subsp. *vulgaris* Hildenborough encodes a potential transmembrane redox protein complex. J Bacteriol 1993;175(15):4699–4711.

42. Dolla A, Pohorelic BK, Voordouw JK, Voordouw G. Deletion of the *hmc* operon of *Desulfovibrio vulgaris* subsp. *vulgaris* Hildenborough hampers hydrogen metabolism and low-redox-potential niche establishment. Arch Microbiol 2000;174(3):143–151.

43. Berks BC, Palmer T, Sargent F. The Tat protein translocation pathway and its role in microbial physiology. Adv Microb Physiol 2003;47:187–254.

44. Kelley LA, Mezulis S, Yates CM, Wass MN, Sternberg MJE. The Phyre2 web portal for protein modeling, prediction and analysis. Nature Protocols 2015;10(6):845–858.

45. Dave JA, Gey van Pittius NC, Beyers AD, Ehlers MR, Brown GD. Mycosin-1, a subtilisin-like serine protease of *Mycobacterium tuberculosis*, is cell wall-associated and expressed during infection of macrophages. BMC Microbiol 2002;2:30.

46. Bunduc CM, Fahrenkamp D, Wald J, Ummels R, Bitter W, et al. Structure and dynamics of a mycobacterial type VII secretion system. Nature 2021;593(7859):445-448.

47. Rawlings ND, Barrett AJ. Evolutionary families of peptidases. Biochem J 1993;290 (Pt 1)(Pt 1):205-218.

48. Joshi MV, Mann SG, Antelmann H, Widdick DA, Fyans JK, et al. The twin arginine protein transport pathway exports multiple virulence proteins in the plant pathogen *Streptomyces scabies*. Mol Microbiol 2010;77(1):252–271.

49. Widdick DA, Dilks K, Chandra G, Bottrill A, Naldrett M, et al. The twin-arginine translocation pathway is a major route of protein export in *Streptomyces coelicolor*. Proc Natl Acad Sci U S A 2006;103(47):17927–17932.

50. Widdick DA, Hicks MG, Thompson BJ, Tschumi A, Chandra G, et al. Dissecting the complete lipoprotein biogenesis pathway in *Streptomyces scabies*. Mol Microbiol 2011;80(5):1395–1412.

51. Schaerlaekens K, Van Mellaert L, Lammertyn E, Geukens N, Anne J. The importance of the Tat-dependent protein secretion pathway in *Streptomyces* as revealed by phenotypic changes in *tat* deletion mutants and genome analysis. Microbiology 2004;150(Pt 1):21–31.

52. Egan AJF, Errington J, Vollmer W. Regulation of peptidoglycan synthesis and remodelling. Nat Rev Microbiol 2020;18(8):446–460.

53. Keenan T, Dowle A, Bates R, Smith MCM. Characterization of the *Streptomyces coelicolor* glycoproteome reveals glycoproteins important for cell wall biogenesis. mBio 2019;10(3).

54. Mazmanian SK, Liu G, Ton-That H, Schneewind O. *Staphylococcus aureus* sortase, an enzyme that anchors surface proteins to the cell wall. Science 1999;285(5428):760-763.

55. Spirig T, Weiner EM, Clubb RT. Sortase enzymes in Gram-positive bacteria. Mol Microbiol 2011;82(5):1044–1059.

56. Schneewind O, Model P, Fischetti VA. Sorting of protein A to the staphylococcal cell wall. Cell 1992;70(2):267–281.

57. Biswas L, Biswas R, Nerz C, Ohlsen K, Schlag M, et al. Role of the twin-arginine translocation pathway in *Staphylococcus*. J Bacteriol 2009;191(19):5921–5929.

58. Malik A, Kim SB. A comprehensive *in silico* analysis of sortase superfamily. J Microbiol 2019;57(6):431–443.

59. Duong A, Capstick DS, Di Berardo C, Findlay KC, Hesketh A, et al. Aerial development in *Streptomyces coelicolor* requires sortase activity. Mol Microbiol 2012;83(5):992–1005.

60. Ize B, Stanley NR, Buchanan G, Palmer T. Role of the *Escherichia coli* Tat pathway in outer membrane integrity. Mol Microbiol 2003;48(5):1183–1193.

61. Bernhardt TG, de Boer PA. The *Escherichia coli* amidase AmiC is a periplasmic septal ring component exported via the twin-arginine transport pathway. Mol Microbiol 2003;48(5):1171–1182.

62. Thompson BJ, Widdick DA, Hicks MG, Chandra G, Sutcliffe IC, et al. Investigating lipoprotein biogenesis and function in the model Gram-positive bacterium *Streptomyces coelicolor*. Mol Microbiol 2010;77:943–955.

63. Widdick DA, Eijlander RT, van Dijl JM, Kuipers OP, Palmer T. A facile reporter system for the experimental identification of twin-arginine translocation (Tat) signal peptides from all kingdoms of life. J Mol Biol 2008;375(3):595–603.

64. Craney A, Tahlan K, Andrews D, Nodwell J. Bacterial transmembrane proteins that lack N-terminal signal sequences. PLoS One 2011;6(5):e19421.

65. Lin Z, Akin H, Rao R, Hie B, Zhu Z, L, et al. Evolutionary-scale prediction of atomic-level protein structure with a language model. Science 2023;379(6637):1123-1130.

66. Keon RG, Fu R, Voordouw G. Deletion of two downstream genes alters expression of the *hmc* operon of *Desulfovibrio vulgaris* subsp. *vulgaris* Hildenborough. Arch Microbiol 1997;167(6):376–383.

